# The potential role of human immune cells in the systemic dissemination of enterovirus-D68

**DOI:** 10.1101/2022.12.01.518644

**Authors:** Brigitta M. Laksono, Syriam Sooksawasdi Na Ayudhya, Muriel Aguilar-Bretones, Carmen W. E. Embregts, Gijsbert P van Nierop, Debby van Riel

**Affiliations:** Department of Viroscience, Erasmus MC, Rotterdam, South Holland, the Netherlands

## Abstract

Enterovirus-D68 (EV-D68) often causes mild respiratory infections, but can also cause severe respiratory infections and extra-respiratory complications, including acute flaccid myelitis (AFM). Systemic dissemination of EV-D68 is crucial for the development of extra-respiratory diseases, but it is currently unclear how EV-D68 viremia occurs. We hypothesize that immune cells contribute to the systemic dissemination of EV-D68, as this is a mechanism commonly used by other enteroviruses. Therefore, we investigated the susceptibility and permissiveness of human primary immune cells for different EV-D68 isolates. In human peripheral blood mononuclear cells (PBMC) inoculated with EV-D68, only B cells were susceptible but virus replication was limited. However, B cell-rich cultures, such as Epstein-Barr virus-transformed B-lymphoblastoid cell line (BLCL) and primary lentivirus-transduced B cells, were productively infected. In BLCL, neuraminidase treatment to remove α2,6- and α2,3-linked sialic acids resulted in a significant decrease of EV-D68 infected cells, suggesting that sialic acids are the functional receptor on B cells. Subsequently, we showed that dendritic cells (DCs), particularly immature DCs, are susceptible and permissive for EV-D68 infection and that they can spread EV-D68 to autologous BLCL. Altogether, our findings suggest that immune cells, especially B cells and DCs, play an important role in the development the systemic dissemination of EV-D68 during an infection, which is an essential step towards the development of extra-respiratory complications.

**Author summary:** Enterovirus D68 (EV-D68) is an emerging respiratory virus that has caused outbreaks worldwide since 2014. EV-D68 infects primarily respiratory epithelial cells and the infection commonly results in mild respiratory diseases. However, EV-D68 infection is also associated with complications outside the respiratory tract, including a polio-like paralysis. Despite the severity of these extra-respiratory complications, it is unclear how EV-D68 is able to spread outside the respiratory tract and infect other organs, like the central nervous system (CNS). To understand this, we investigated if immune cells play a role in the extra-respiratory spread of EV-D68. We showed that EV-D68 can infect and replicate in specific immune cells, *i*.*e*. B cells and dendritic cells (DCs), and that the virus can be transferred from DCs to B cells. Our findings suggest that lymphoid tissues, which harbor many immune cells, can be a secondary replication site for EV-D68, from where virus is released in the circulation. Our data reveal the importance of immune cells in the systemic spread of EV-D68, which is essential for infection of extra-respiratory tissues. Intervention strategies that prevent EV-D68 infection of immune cells will therefore potentially prevent virus spread from the respiratory tract to other organs.

## Introduction

Enterovirus D68 (EV-D68) is a small non-enveloped, positive single-stranded RNA virus that belongs to the family *Picornaviridae*, genus *Enterovirus*. Although EV-D68 causes predominantly mild upper respiratory tract symptoms, EV-D68 caused outbreaks of severe respiratory diseases worldwide in 2014, which were associated with neurological complications in some individuals. Among these complications, acute flaccid myelitis (AFM) was most frequently reported (1-4). Since then, EV-D68 has caused biennial outbreaks of severe respiratory disease and AFM up to 2018 (3, 5). EV-D68 circulation was limited during the pandemic, partly due to COVID-19 measures, but EV-D68-associated severe respiratory illnesses have been rising in several countries since 2021 and 2022 due to easing of COVID-19 measures (6, 7). Throughout the years, multiple clades of EV-D68 circulated, but so far there is little evidence for differences in the virulence of these viruses (1, 8, 9).

The ability of EV-D68 to disseminate into the circulation (viremia) is essential for the virus to spread from the respiratory tract, which is the primary replication site of the virus, to other organs, such as the central nervous system (CNS), and cause extra-respiratory complications. However, despite the importance of viremia in the pathogenesis of EV-D68 infection, the mechanism that leads to EV-D68 viremia is poorly understood.

Studies in mice, ferrets and patient samples have shown that the virus or viral RNA can be detected in the circulation and extra-respiratory tissues (1, 10-17). In intranasally inoculated mice, virus was detected in the blood within 24 hours post-inoculation (hpi), and in extra-respiratory tissues, such as the spleen and skeletal muscles (10, 11). In intranasally inoculated ferrets, virus was detected in axillary lymph nodes at multiple days post-inoculation (earliest detection at day 5 post-inoculation) (12). In humans, virus and viral RNA have been detected in serum of EV-D68 patients, but it is currently unknown how frequent viremia occurs during an EV-D68 infection (13-17). In addition, it is unclear whether the virus detected in the circulation is a direct spill-over from the respiratory tract or whether virus first spreads to and replicates in other tissues, *e*.*g*. lymphoid tissues, before disseminating into the circulation.

Other EVs, such as poliovirus and EV-A71, are known to replicate in lymphoid tissues, resulting in a sustained production of infectious viruses and subsequent spill-over into the circulation (18-22). In the case of EV-D68, the role of immune cells and lymphoid tissues in the development of viremia remains unclear. Previous studies have shown that EV-D68 productively infects several human immune cell lines, such as granulocytic (KG-1), monocytic (U-937), T (Jurkat and MOLT) and B (Raji) cell lines (23), suggesting that immune cells or lymphoid tissues may play a role in the development of EV-D68 viremia. Here, we investigated the susceptibility and permissiveness of human primary immune cells to infection of EV-D68 from different clades to unravel the potential role of immune cells in the development of viremia and to investigate if there are clade-specific differences in the susceptibility and permissiveness of these cells. Subsequently, we investigated whether dendritic cells (DCs) can transmit virus to other immune cells, such as B cells.

## Results

### Human B cells are susceptible and permissive to EV-D68 infection

To investigate whether human leukocyte subsets are susceptible to EV-D68 infection, human peripheral blood mononuclear cells (PBMC) were inoculated with EV-D68 strains from subclades A, B2 and D. We observed that B cells were susceptible to infection with EV-D68/A (mean 6% ± standard error of mean (SEM) 1) and EV-D68/B2 (3% ± 1) but not for EV-D68/D, based on intracellular expression of EV-D68 capsid protein VP1. In one donor, 3% of monocytes were susceptible to infection with EV-D68/A. We did not observe any VP1^+^ cells in CD4^+^ nor CD8^+^ T cell population. (**Fig 1A**).

**Fig 1.**
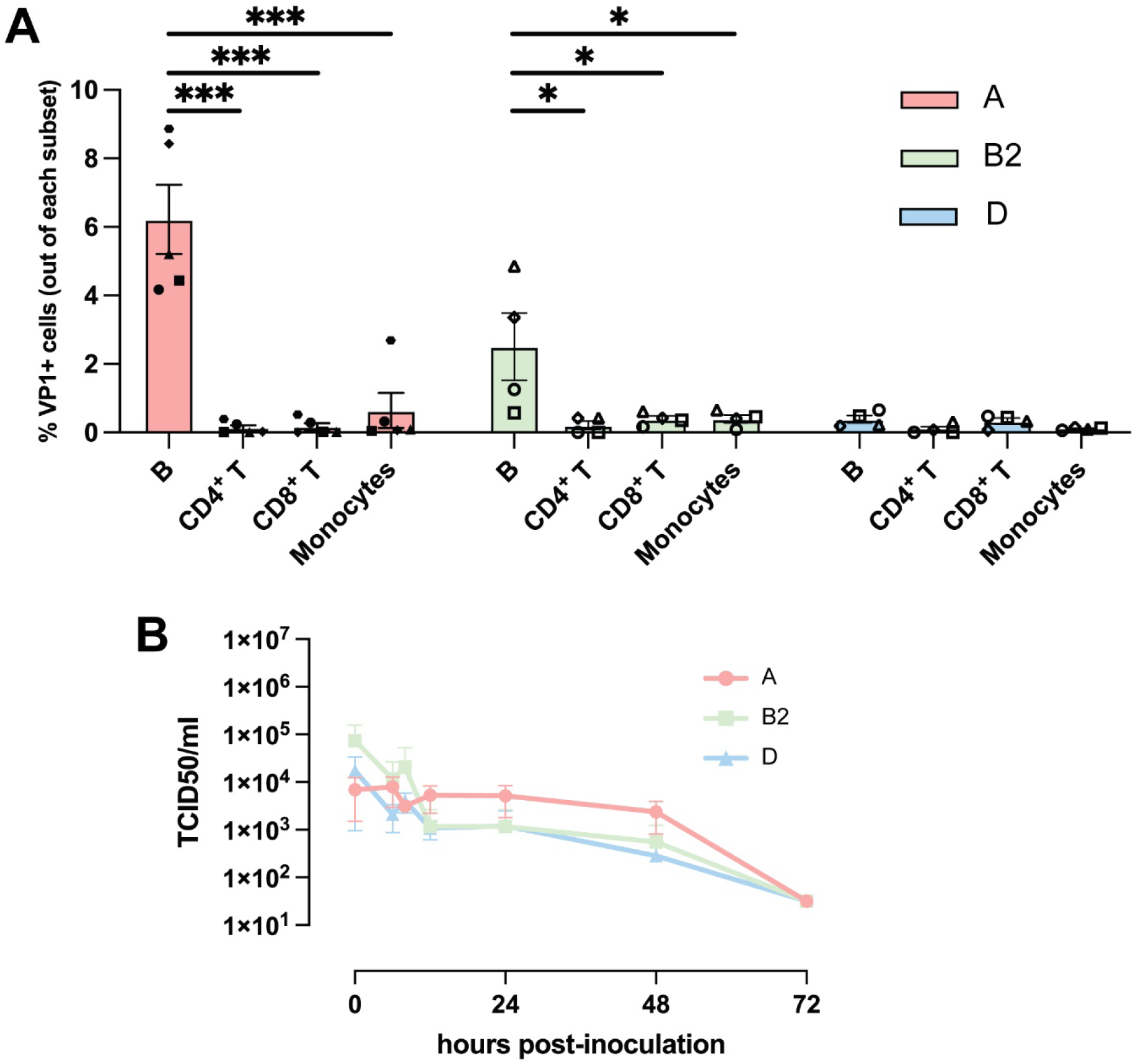
Susceptibility and permissiveness of human PBMC to EV-D68 infection. (A) Percentage of EV-D68 VP1^+^ leukocyte subsets 24 h after inoculation with EV-D68 strains from subclades A (n = 5 donors), B2 (n = 4 donors) and D (n = 4 donors). Each symbol represents one donor. Statistical analysis was performed using a one-way ANOVA with multiple comparison test. Error bars denoted SEM. (B) Production of infectious viruses in EV-D68-inoculated PBMC. PBMC: peripheral blood mononuclear cells; SEM: standard error of mean *: P<0.05; ***: P≤0.001.

Next, we investigated whether EV-D68 inoculation of PBMC resulted in a productive infection. Supernatant and cell lysate were collected and the presence of infectious virus particles at 0, 6, 8, 10, 24, 48 and 72 hpi was determined by virus titration. Despite the presence of infected cells after inoculation with EV-D68/A or EV-D68/B2, we did not observe an increase in infectious virus titer at any time point (**Fig 1B**). As human PBMC consists only of 4 – 14% of B cells, (24) of which 3 – 6% were infected by EV-D68, we cannot exclude the possibility that viral replication occurs in VP1^+^ B cells but this was below the detection limit of the assay.

In order to determine whether B cells are susceptible and permissive for EV-D68, we utilized two B cell-rich models. We first inoculated Epstein-Barr virus-transformed B-lymphoblastoid cell line (BLCL) with EV-D68 strains from subclades A, B2 and D. The inoculation resulted in 13% ± 4 EV-D68/A, 18% ± 5 EV-D68/B2 and 21% ± 4 EV-D68/D VP1^+^ cells, although there was variation among donors in the percentage of infected cells (**Fig 2A**). Production of new infectious viruses over time (∼2 – 3 logarithmic increase of TCID_50_/ml within 24 hours) was detected after inoculation with all three viruses, in which EV-D68/B2 and EV-D68/D replicated faster than EV-D68/A, albeit without any statistical differences (**Fig 2B**).

**Fig 2.**
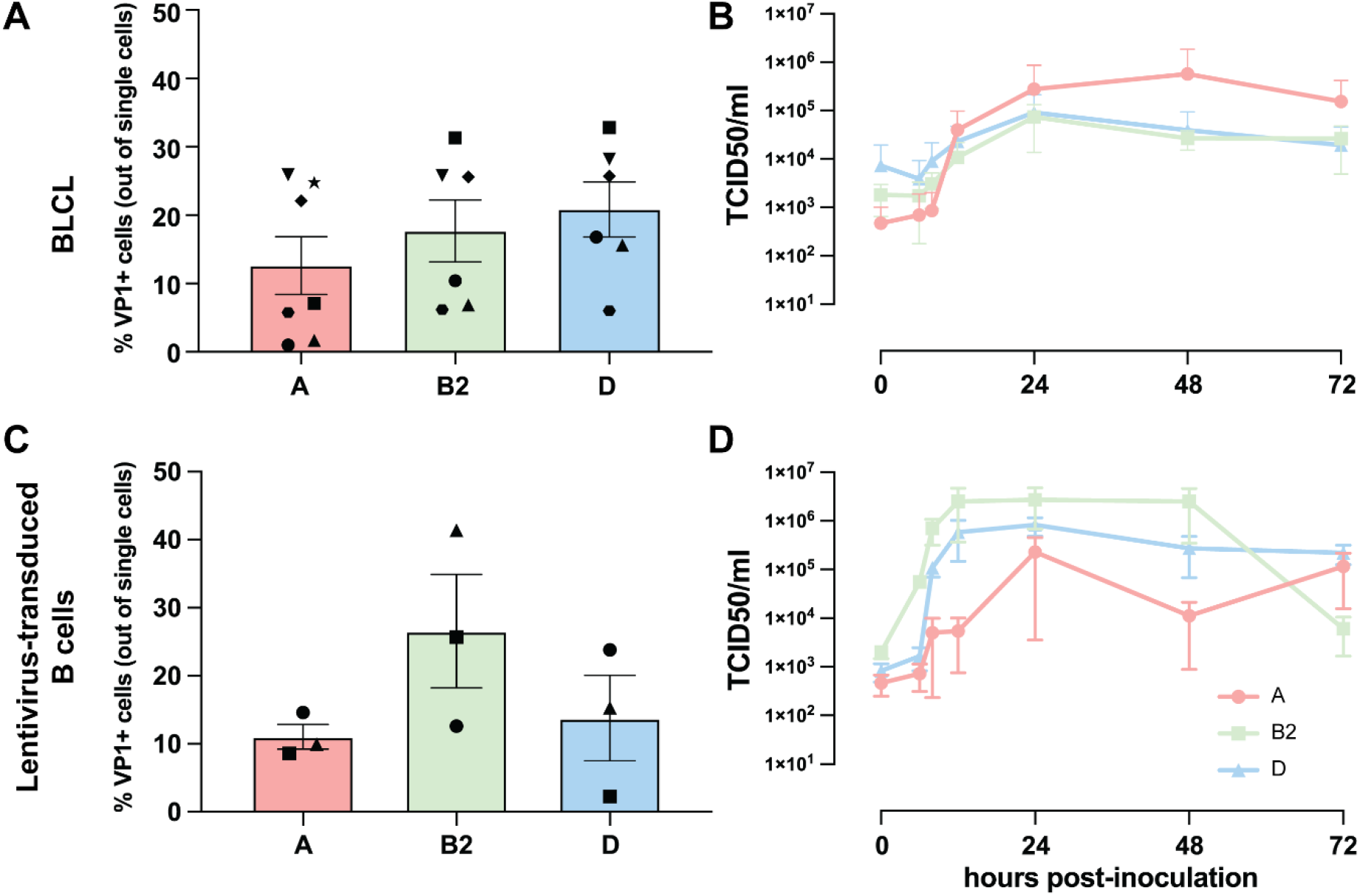
Susceptibility and permissiveness of B cell-rich models to EV-D68 infection. BLCL and lentivirus-transduced B cells were inoculated with EV-D68 strains from subclades A (n = 7), B2 (n = 6) and D (n = 6) as models for EV-D68 infection in B cell-rich environment. (A – B) Percentage of EV-D68 VP1^+^ cells at 24 hpi and production of new infectious virus particles in EV-D68-inoculated BLCL over time, respectively. (C – D) Percentage of VP1^+^ cells at 24 hpi and production of infectious viruses in EV-D68-inoculated lentivirus-transduced B-cells. Each symbol in (A) and (C) represents one donor. No statistically significant differences were observed in the percentages of VP1^+^ cells among the different virus clades. Statistical analysis was performed using a one-way ANOVA with multiple comparison test. Error bars denote SEM. BLCL: B-lymphoblastoid cell line; hpi: hours post-inoculation; SEM: standard error of mean.

Primary B cell clones that were lentivirus-transduced to express the germinal center-associated B cell lymphoma-6 (Bcl-6) and Bcl-xL in order to endow a stable proliferative state, were inoculated with EV-D68 strains from subclades A, B2 and D. This resulted in 11% ± 2 EV-D68/A, 27% ± 8 EV-D68/B2 and 14% ± 6 EV-D68/D VP1^+^ cells (**Fig 2C)**. The inoculation also resulted in production of new infectious viruses (∼2 – 3 logarithmic increase within 10 to 24 hours), without any statistical differences among the different EV-D68 clades **(Fig 2D)**.

### Infection of BLCL is largely mediated by the presence of α2,3- and α2,6-linked SAs

Several immune cells, including B cells, express α2,6-linked and α2,3-linked sialic acids (SAs), which can function as receptors for EV-D68 to initiate binding and virus entry (25-27). To investigate whether α2,3- and α2,6-linked SAs mediate EV-D68 infection of BLCL, BLCL were treated with *Arthrobacter ureafaciens* neuraminidase (ANA) to remove cell surface SAs prior to inoculation with EV-D68 strains. Upon ANA treatment, the average percentages of α2,3-linked SA^+^ BLCL decreased from 77% ± 9 to 38% ± 3 and of α2,6-linked SA^+^ cells from 89% ± 1 to 5% ± 1 (**Fig 3A**). ANA treatment of BLCL prior to inoculation resulted in a significant decrease of percentage of VP1^+^ cells, with only 2% ± 1 EV-D68/A, 5% ± 2 EV-D68/B2 and 5% ± 1 EV-D68/D VP1^+^ cells (**Fig 3B**).

**Fig 3.**
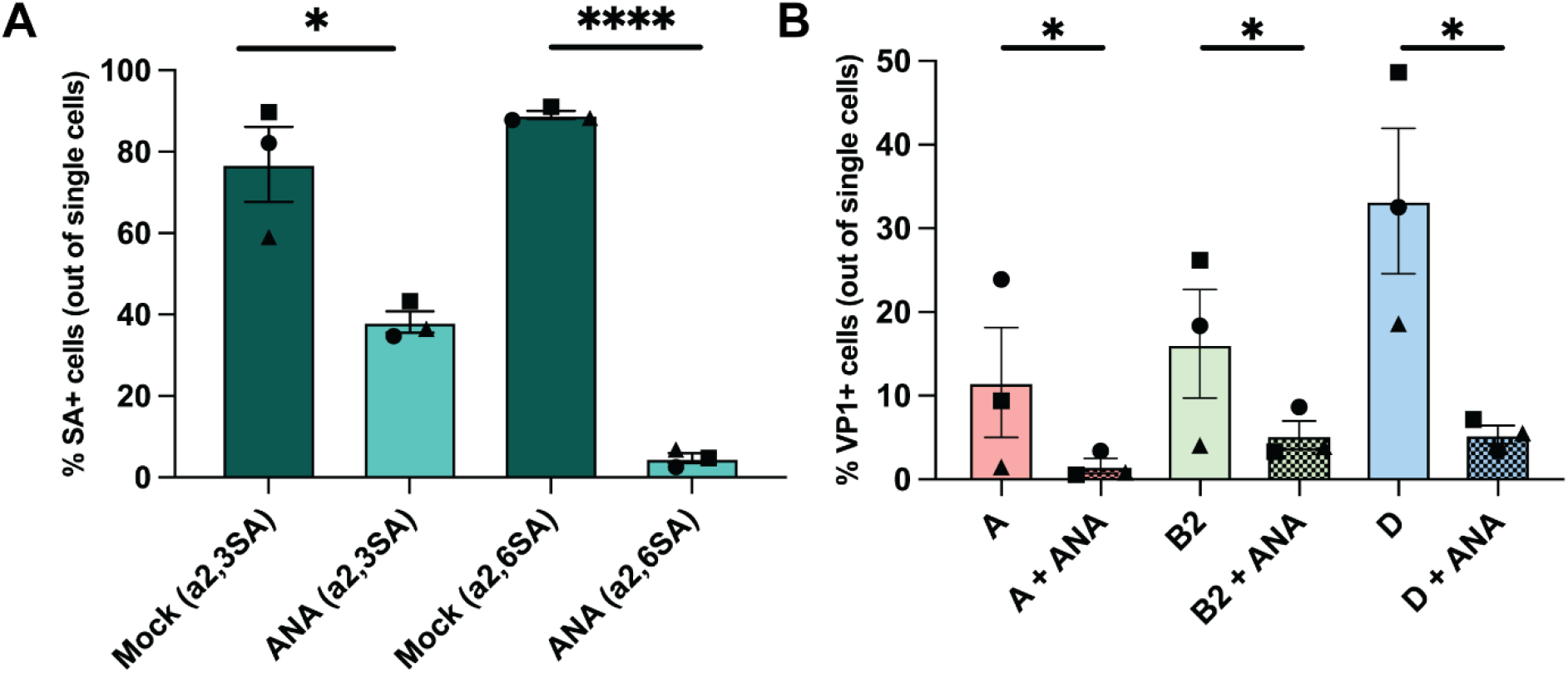
Percentages of α2,3-and α2,6-linked SAs^+^ and EV-D68 VP1^+^ BLCL upon neuraminidase treatment. (A) Percentage of BLCL expressed α2,3- (n=3) and α2,6- (n=3) linked SAs with and without ANA treatment. (B) Percentage of VP1^+^ BLCL (n=3) with and without ANA treatment inoculated with EV-D68 from subclades A, B2 and D, measured 24 hpi. Statistical analysis was performed with t-test. Error bars denote SEM. BLCL: B-lymphoblastoid cell line. ANA: *Arthrobacter ureafaciens* neuraminidase; SAs: sialic acids; hpi: hours post-inoculation; SEM: standard error of mean. *: P<0.05; ****: P≤0.0001.

### Dendritic cells (DCs) are susceptible and permissive to EV-D68 infection

DCs are a subset of immune cells that are attracted to sites of inflammation, but not abundantly present in PBMC (28). To investigate whether DCs are susceptible and permissive to EV-D68 infection, monocytes were differentiated in immature and mature DCs (imDCs and mDCs, respectively), and inoculated with EV-D68 strain from subclade A as a representative strain. Increased expression of maturation markers (HLA-DR, CD86, PD-L1 and CD83) was used to confirm differentiation of imDCs to mDCs (**S1 Fig)**. The average percentage of VP1^+^ cells in imDCs (10% ± 1) was significantly higher than in mDCs (4% ± 1) at 6 hpi (**Fig 4A**). From 2 to 10 hpi, viral titers in the supernatants increased ∼1 logarithmic TCID_50_/ml in imDCs and ∼0.5 logarithmic TCID_50_/ml in mDCs inoculated with EV-D68/A, after which the virus titers decreased. Despite this decrease, at 48 hpi, virus titer in the supernatants of imDCs were significantly higher than in mDCs (**Fig 4B**).

**Fig 4.**
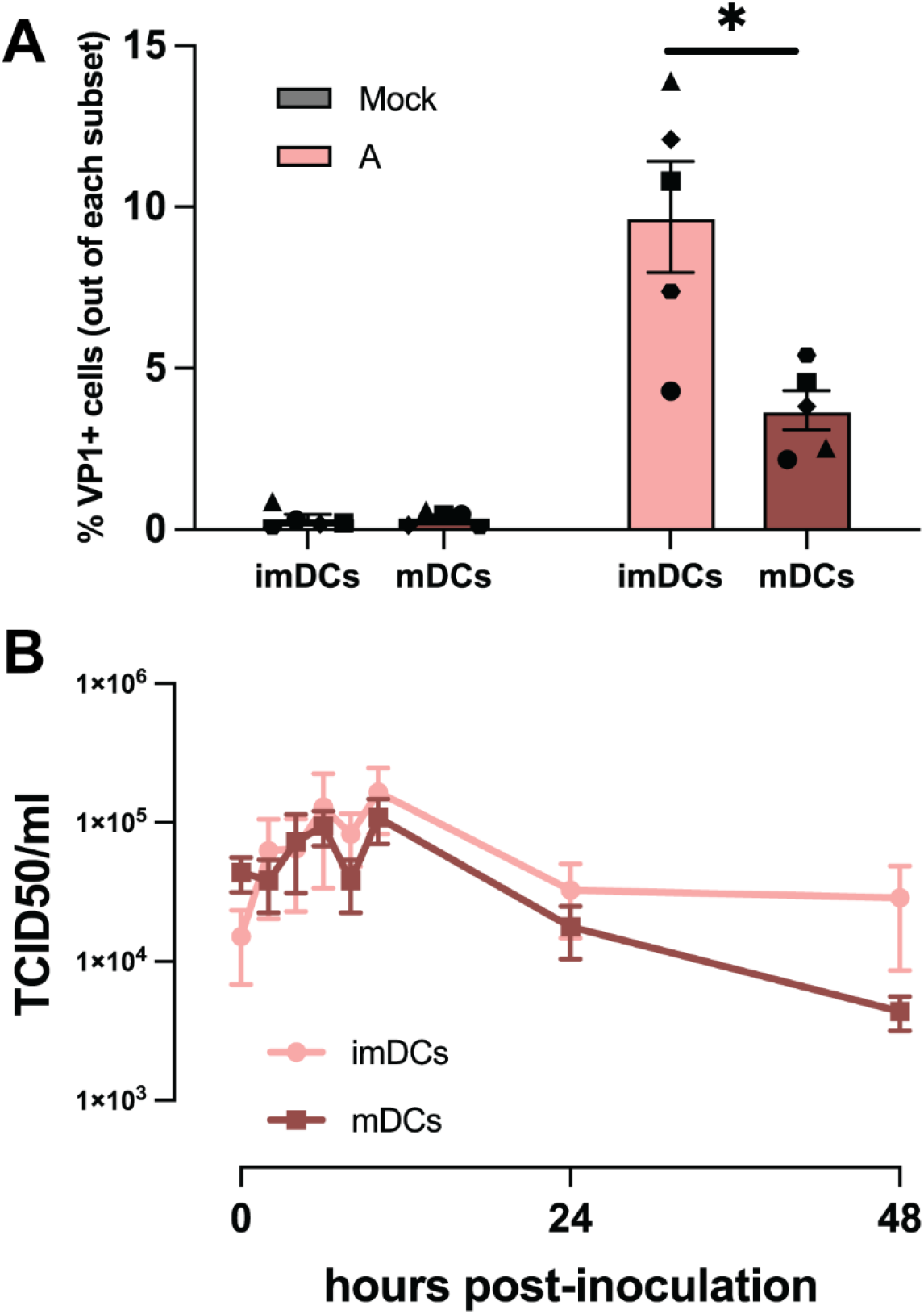
Susceptibility and permissiveness of imDCs and mDCs to EV-D68 infection. (A) Percentage of EV-D68 VP1^+^ imDCs (n = 5) and mDCs (n = 5) measured at 6 hpi. (B) Production of infectious viruses in EV-D68/A-inoculated imDCs (n = 3) and mDCs (n = 3). Samples for virus titration were collected at 0, 2, 4, 6, 8, 10, 24 and 48 hpi. All statistical analyses in this figure were performed with t-test. Error bars denote SEM. imDCs: immature dendritic cells; mDCs: mature dendritic cells; hpi: hours post-inoculation; SEM: standard error of mean. *: P<0.05; **: P<0.01.

### imDCs can tranfer EV-D68 to autologous BLCL

Dendritic cells are important for local antiviral responses and antigen take-up induces DC maturation and migration to lymphoid tissues (28). Therefore, we investigated whether imDCs can transmit EV-D68 infection to B cells. EV-D68/A-inoculated imDCs were co-cultured with autologous BLCL (BLCL+DC), after which the percentages of VP1^+^ BLCLs were determined. The following controls were included: (1) EV-D68/A-inoculated imDCs (DC only); (2) BLCL cultured with supernatant from EV-D68/A-inoculated imDCs collected at 0 hpi, directly after the virus inoculation for 1 hour and the subsequent washing steps (BLCL + t0 DC sup) and (3) BLCL cultured with supernatant from EV-D68/A-inoculated imDCs collected at 6 hpi (BLCL + t6 DC sup). The schematic representation of this experiment is presented in **Fig 5A**. As observed previously, inoculation of imDCs in the absence of their autologous BLCL resulted in an average of 11% ± 2 VP1^+^ cells at 6 hpi and the percentage remained stable at 24 hpi. In imDCs co-cultured with BLCL, we observed an average percentage of 10% ± 4 VP1^+^ imDCs, although this percentage decreased at 24 hpi (4% ± 2) (**Fig 5B**). When BLCLs were inoculated with supernatants from EV-D68-inoculated imDCs collected at 0 or 6 hpi, only very few B cells were infected, with average percentages of 0.5% VP1^+^ BLCLs in BLCL+t0 DC sup and 1% VP1^+^ BLCLs in BLCL+t6 DC sup at 24 hpi. When BLCLs were co-cultured with EV-D68-inoculated imDCs, the percentage of infected BLCL increased from 2%±1 at 6 hpi to 6% ± 1 at 24 hpi (**Fig 5C**).

**Fig 5.**
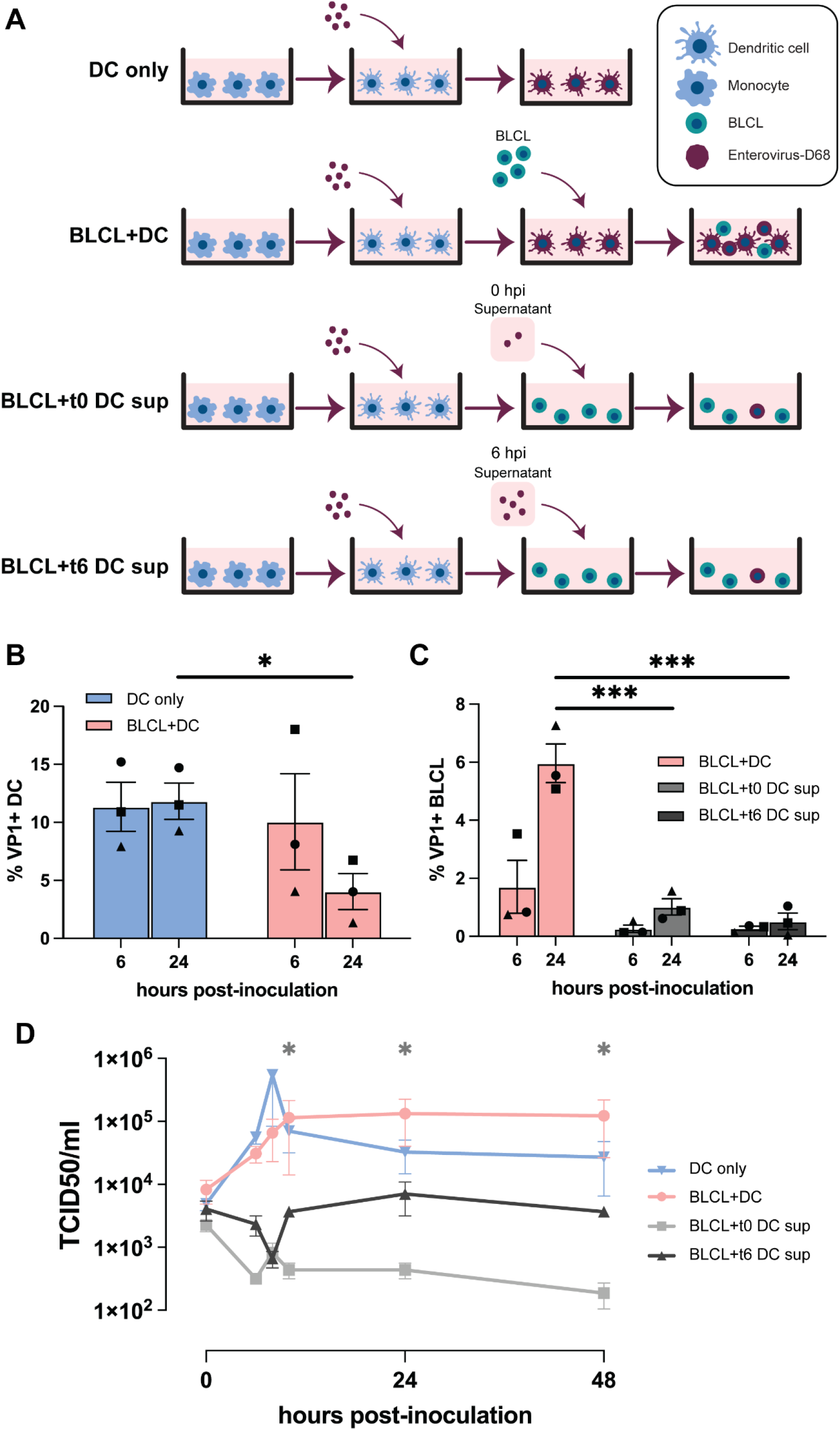
Co-culture of EV-D68-inoculated imDCs with their autologous BLCL. (A) Schematic representation of the experimental setup and controls. (B) Percentages of EV-D68 VP1^+^ imDCs and (C) BLCL in different culture conditions at 6 and 24 hpi. (D) Production infectious viruses in different culture conditions. Statistical analysis in (B) was performed with t-test. Statistical analysis in (C) and (D) were performed with a one-way ANOVA with multiple comparison test and compared to BLCL+t0 DC sup and BLCL+t6 sup, respectively. Error bars denote SEM. BLCL: B-lymphoblastoid cell line; imDCs: immature dendritic cells; hpi: hours post-inoculation; SEM: standard error of mean; BLCL+t0 DC sup: autologous BLCL co-cultured with supernatant collected from EV-D68/A-inoculated imDCs at 0 hpi. BLCL+t6 DC sup: autologous BLCL co-cultured with supernatant collected from EV-D68/A-inoculated imDCs at 6 hpi. *: P<0.05; ***: P≤0.001.

The inoculation of imDCs in the absence of BLCL resulted in production of new infectious virus particles (∼1.5 logarithmic increase of TCID_50_/ml) within 24 h. Detection of infectious virus particles in the supernatants of the BLCL+DC co-culture showed efficient virus replication and stable virus titer over time (∼1 logarithmic increase of TCID_50_/ml) within the same period of time. Very few, if any, new infectious virus particles were produced by BLCL inoculated with t0 or t6 DC supernatants (**Fig 5D**). Together, the results suggested that imDCs are capable to transmit virus to other susceptible immune cells, such as B cells, which subsequently become productively infected.

## Discussion

The systemic dissemination of EV-D68 is an essential step for extra-respiratory spread of the virus and the development of associated complications, such as AFM, but the underlying mechanism of how viruses spread to the circulation or the origin of virus in blood is largely unknown. In this study, we reveal the potential role of immune cells in the systemic dissemination of EV-D68. We show that human B cells and DCs are susceptible and permissive to EV-D68 infection and that DCs may play a role in the transmission of virus to B cells.

Immune cells are susceptible to infection of other members of *Picornaviridae*. Coxsackievirus type B infects murine splenic B cells resulting in production of new virus particles (29). Poliovirus productively infects DCs and macrophages *in vitro* (human PBMC) and *in vivo* (non-human primates) (30, 31). Echoviruses and EV-A71 have also been reported to infect human imDCs and mDCs (32, 33). Here, we showed that primary human B cells and DCs are susceptible and permissive to EV-D68 infection, which fits with findings in a cohort study that detected enterovirus RNA in peripheral blood B cells and DCs (34). In B cells, this tropism is facilitated by the presence of α2,6- and/or α2,3-linked SAs. Productive infection was only detected in BLCLs and lentivirus-transduced B cells; the latter resembles activated germinal center B cells (35). Additionally, imDCs were more susceptible and permissive to EV-D68 infection than mDCs. Therefore, it is possible that activation or maturation state of immune cells play a role in the susceptibility and permissiveness to EV-D68 infection, similar to what is observed in poliovirus (30). Furthermore, it may be that the different susceptibility to EV-D68 between imDCs and mDCs is due to higher expression of α2,6-linked SA on imDCs than mDCs (36).

We observed differences in the susceptibility and permissiveness to EV-D68 infection among viruses or donors included in this study. Differences in the susceptibility for different EV-D68 clades were observed in B cells within the PBMCs, but this was not observed in BLCLs and lentivirus-transduced B cells. Donor variation was observed in PBMCs and all B cell models and, in one PBMC donor, we observed monocytes infected with EV-D68 isolate from subclade A. The underlying mechanism for these differences among donors and possibly among different virus clades, as well as their association with the risk of the development of extra-respiratory diseases, are still unclear.

Virus replication within immune cells and/or lymphoid tissues is likely important for the development of a viremia. Due to the susceptibility and permissiveness of immune cells to EV-D68 infection, an immune cell-rich environment, such as a lymphoid tissue, can serve as a secondary replication site for EV-D68, from where the virus can spread into the circulation. Since poliovirus viremia is essential for virus spread to the CNS and the subsequent development of CNS diseases (37), it can be speculated that EV-D68 viremia is essential for virus entry into the CNS and subsequent neurological complications, such as AFM (4, 13). In addition, a viremia could lead to virus spread to other organ system contributing to non-neurological complications associated with EV-D68 infection, including acute gastroenteritis, myocarditis and skin rash (38-40). Prevention of spread or productive infection in the lymphoid tissues may prevent systemic dissemination, as observed in poliovirus infection, in which vaccination prevents viral spread to other organs (41).

Based on our findings, we propose a model that explains the systemic dissemination of EV-D68 (**Fig 6**). EV-D68 infects respiratory epithelial cells, which results in the production of infectious virus particles and the recruitment of immune cells, such as imDCs. From the respiratory tract, cell-free virus can spill over into the circulatory and lymphatic systems and spread into lymphoid tissues. Alternatively, imDCs can become infected, and migrate to the lymphoid tissues, where they release newly produced viruses or spread virus to resident DCs or B cells. Lymphoid tissues can be the secondary replication site for EV-D68, from where the virus is released in the bloodstream. Prevention of virus spread to and amplification in lymphoid tissues can therefore prevent the development of a subsequent viremia and severe extra-respiratory complications caused by EV-D68.

**Fig 6.**
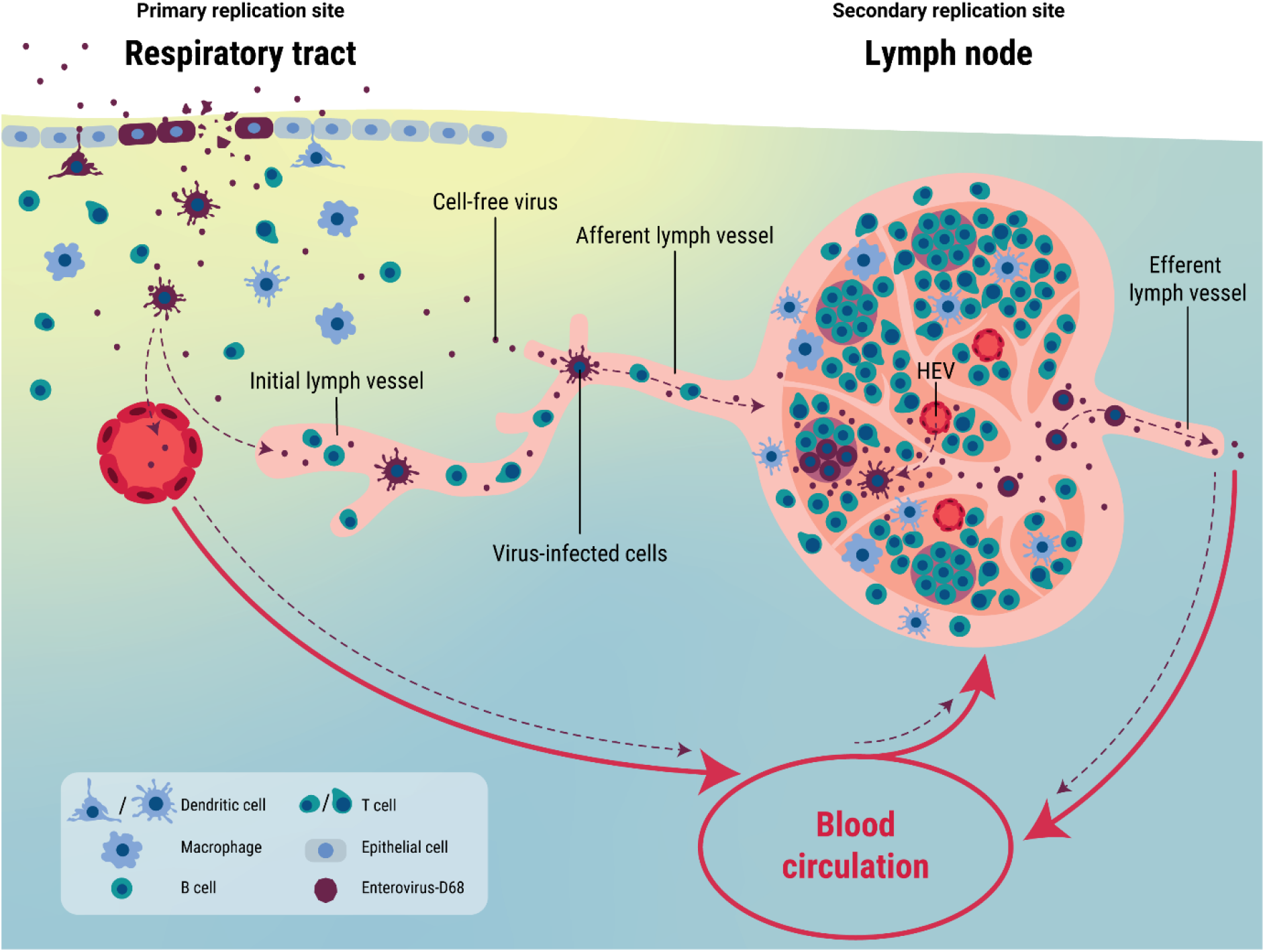
Proposed model for systemic dissemination of EV-D68. EV-D68 enters the respiratory tract and initially infects respiratory epithelial cells. The infection will result in the recruitment of immune cells, including imDCs. Subsequently, cell-free EV-D68 can spread to lymphoid tissues by spilling over into the circulatory or lymphatic system. Alternatively, it can infect imDCs, which can transfer the virus to lymphoid tissues. These lymphoid tissues can function as a secondary replication site, where EV-D68 infects mDCs and B cells. From this site, cell-free virus or virus-infected immune cells can enter the blood circulation and spread virus to other tissues. HEV: high endothelial venule. Red arrow: blood circulation; purple dotted arrow: potential routes of EV-D68 spread.

## Materials and methods

### Ethics statement

PBMC were obtained from healthy adult donors after obtaining written informed consent. The studies were approved by the medical ethical committee of Erasmus MC, the Netherlands (MEC-2015-095). For experiment involving human buffy coats, written informed consent for research use was obtained by the Sanquin blood bank.

### Cells

Human PBMC were isolated from blood (n = 8 healthy donors) by Ficoll density gradient centrifugation. BLCL were established from 5 donors by transformation with Epstein-Barr virus as previously described (42). PBMC and BLCL were cultured in RPMI-1640 medium (Capricorn) supplemented with 10% fetal bovine serum (FBS) and 100 IU/ml of penicillin, 100 µg/ml of streptomycin and 2 mM L-glutamine (PSG). After virus inoculation, supernatants and cell lysates were collected and frozen and thawed three times prior to sample processing.

Germinal center-like primary B cell clones were generated from 3 donors. Synthetic cDNA encoding a self-cleaving polyprotein Bcl-6.t2A.Blc-xL to express the germinal center B cell-associated transcription factors Bcl-6 and Bcl-xL was synthesized (B6L; Integrated DNA Technologies) and cloned in pENTR/D-TOPO (Thermo Fisher), generating pENTR.B6L (35, 43). The B6L cDNA was subsequently transferred to pLenti6.3/V5-DEST using Gateway LR Clonase II (Thermo Fisher), generating pLV-B6L. Subsequently, lentiviral vector stocks (LV-B6L) were generated, all according to manufacturer’s instructions (Thermo Fisher). Primary CD19^+^ B-cells were isolated from human PBMC using the EasySep human CD19 positive selection kit II (StemCell Technologies) and transduced using LV.B6L as described elsewhere (35). LV-B6L transduced B cells were cultured in AIM-V AlbuMAX medium (Gibco) supplemented with 10% FBS, PSG, 50 µM beta-mercaptoethanol (Sigma), 25 ng/ml recombinant human IL-21 (Peprotech) and growth-arrested 40 gray gamma-irradiated L-CD40L feeder cells. Every 3 to 4 days, culture medium was refreshed with 25 ng/ml recombinant human IL-21 and new growth-arrested 40 gray gamma-irradiated L-CD40L feeder cells. Clonal B cell cultures from each donor were generated using limiting dilution. Absence of antibody reactivity towards EV-D68 was confirmed to exclude antibody mediated effects on EV-D68 infection (data not shown). After virus inoculation, supernatants and cell lysates were collected and frozen and thawed three times prior to sample processing.

Human rhabdomyosarcoma (RD) cells were cultured in DMEM (Lonza) supplemented with 10% FBS and PSG.

### Viruses

EV-D68 strains included in this study were isolated from clinical specimens at the National Institute of Public Health and the Environment (RIVM), Bilthoven, the Netherlands. The viruses were isolated on RD cells (ATCC) at 33°C at RIVM from respiratory samples from patients with EV-D68-associated respiratory disease. Virus stocks for the studies were grown in RD cells at 33°C in 5% CO_2_. The viruses included in this study with virus reference number, year of isolation an accession number are as follows: clade A (or A1) (4311200821; 2012, accession number MN954536), clade D (or A2) (4311400720; 2014, accession number MN954537) and subclade B2 (4311201039; 2012, accession number MN954539). All virus stocks were sequenced for their whole genome, and that there was no evidence for cell culture adaptive mutations.

### Virus titration

Virus titers were assessed by end-point titrations in RD cells and were expressed in median tissue culture infectious dose (TCID_50_/ml). In brief, 10-fold serial dilutions of a virus stock were prepared in triplicate and inoculated onto a monolayer of RD cells. The inoculated plates were incubated at 33°C in 5% CO_2_. Cytopathic effect (CPE) was determined at day 5, and virus titers were determined using the Spearman-Kärber method (44).

### Differentiation of monocyte-derived DCs

Human monocytes were isolated from buffy coats (n= 5 donors) by Ficoll density gradient centrifugation and selected for CD14^+^ cells using magnetic beads. Monocytes were cultured in RPMI-1640 medium supplemented with 1XGlutamax, 10% FBS, 100 IU/ml of penicillin and 100 µg/ml of streptomycin. Subsequently, monocytes were differentiated into imDCs in the presence of human interleukin 4 (IL-4; 20ng/ml; PEPROTECH; 200-04) and human granulocyte-macrophage colony-stimulating factor (GM-CSF; 20ng/ml; MILTENYI BIOTEC; 130-093-866). At day 5, monocyte-derived imDCs were further differentiated into mature DCs by adding lipopolysaccharide (LPS; 1ug/ml; Thermo Fisher Scientific; L8274-10MG) into the medium. mDCs were defined by the increased expression of HLA-DR, CD86, PD-L1 and CD83 compared to imDCs (**S1 Fig**) and the determination of the cellular marker expression by flow cytometry is described below.

### EV-D68 infection of PBMC, BLCL and lentivirus-transduced B cells

Freshly isolated PBMCs, BLCLs or lentivirus-transduced B cells (1×10^6^ cells) were inoculated with EV-D68 at multiplicity of infection (MOI) of 0.1 for 1 h at 37°C in 5% CO_2_. The inoculum was removed and cells were seeded in new medium into a U-bottomed 96-well plate. Cells and supernatants were collected at 0, 6, 8, 12, 24, 48 and 72 h post-inoculation (hpi). The collected specimens were frozen and thawed three times to allow release of intracellular virus before further used for virus titration. Cells were also collected at 6, 24, 48 and 72 hpi for detection of intracellular capsid protein VP1 (10 μg/ml; GeneTex; GTX132313) by flow cytometry as described below.

### Removal of cell surface sialic acids on BLCL

BLCLs were incubated with 50 mU/ml *Arthrobacter ureafaciens* neuraminidase (Roche) in serum-free medium for 2 h at 37°C in 5% CO_2_. Removal of α(2,3)-linked and α(2,6)-linked sialic acids on the cell surface was verified by staining with biotinylated *Maackia amurensis* lectin (MAL) I (5 μg/ml; Vector Laboratories; B-1265-1) or fluorescein-labeled *Sambucus nigra* lectin (SNA) (5 μg/ml; EY Laboratories; BA-6802-1), respectively. Biotin was detected using a streptavidin-conjugated AlexaFluor488 (5 μg/ml; Thermo Fisher Scientific; S11223). Virus and mock inoculations in non-enzymatic-treated cells were included as positive and negative infection controls, respectively. Infection of BLCL in different conditions were performed as describe above and intracellular capsid protein VP1 were detected by flow cytometry as described below.

### EV-D68 inoculation of DCs and co-culture assays

ImDCs and mDCs in a flat-bottomed 96-well plate (1×10^5^ cells/well) were inoculated with EV-D68/A at MOI of 1. After 1 h, the inoculum was removed and the monocytes-derived imDCs were supplemented with complete RPMI-1640 containing human IL-4 and GM-CSF, while mDCs were supplemented with complete RPMI-1640 containing human IL-4, GM-CSF and LPS. Cells and supernatants were collected at 0, 2, 4, 6, 8, 10, 24 and 48 hpi for virus titration or at 6 hpi for detection of intracellular expression of capsid protein VP1 by flow cytometry as described below.

For co-culture assay, infected imDCs (1×10^5^ cells/well) as describe above were incubated with (2×10^5^ BLCL/well; BLCL+DC) or without autologous BLCL (DC only). To investigate whether imDCs transfer virus particles directly to autologous BLCL or indirectly by release of infectious virus, the supernatant from imDCs that were infected with EV-D68/A for 6 hours, were transferred to autologous BLCL (BLCL+t6 DC sup). As a control, to show that infected BLCL is not due to the leftover of virus inoculum, supernatant from the last washing step in infected imDCs were transferred to the autologous BLCL (BLCL+t0 DC sup). The schematic representation of the co-culture assay is presented in **Fig 5A**. The intracellular expression of capsid protein VP1 was detected at 6 and 24 hpi by flow cytometry as described below. Cells and supernatant were collected to detect infectious virus particles at different time points.

### Flow cytometry

For determination of leukocyte phenotypes, human PBMC were incubated with monoclonal antibodies against CD19 (PE-Cy7; Beckman Coulter; IM3628), CD3 (PerCP; BD Biosciences; 345766), CD4 (V450; BD Biosciences; 560811), CD8 (AmCyan; BD Biosciences; 339188), CD14 (BV711; BD Biosciences; 740773) and CD16 (AlexaFluor647; BD Biosciences; 302020). For determination of DC phenotypes, DCs were incubated with monoclonal antibodies against HLA-DR (Pacific Blue; BioLegend; 307624), CD83(PE-Cy7; BioLegend; 305326), CD86 (AF647; BioLegend; 305416) and PD-L1 (BV785; BioLegend; 320736). For determination of cell phenotypes in DC-BLCL co-cultures, the cells were incubated with monoclonal antibodies against HLA-DR (Pacific Blue; BioLegend; 307624), CD86 (AF647; BioLegend; 305416), CD19 (PE-Cy7; Beckman Coulter; IM3628). Cells were fixed and permeabilized with BD Cytofix/Cytoperm Fixation and Permeabilization kit according to the manufacturer’s instructions (BD Biosciences). The presence of intracellular capsid protein VP1 was determined by staining with polyclonal rabbit anti-EV-D68 VP1 (10 µg/ml; Genetex; GTX132313) and goat anti-rabbit IgG (FITC; BD Biosciences; 554020). Flow cytometry was performed using BD FACS Lyric (BD Biosciences, USA). Data were acquired with BD Suite software and analyzed with FlowJo software. Gating strategies to define different cell phenotypes and to define VP1^+^ cells are presented in **S2 – 5 Figs**.

### Statistical analyses

Statistical analyses were performed using GraphPad Prism 9.0 software (La Jolla, CA, USA). Specific tests are described in the figure legends. P values of ≤0.05 were considered significant. All data were expressed as standard error of mean (SEM).

## Funding

This work was funded by a fellowship to DvR from the Netherlands Organization for Scientific Research (VIDI contract 91718308) and the Erasmus MC Foundation. SSNA received the Royal Thai Government Scholarship supported by the Ministry of Science and Technology of Thailand to perform her doctoral study. GPvN received financial supports from the European Union’s Horizon 2020 research and innovation program under grant agreements no. 874735 (VEO) and no. 101003589 (RECoVER), and the Dutch Research Council (NWO) project no. 109986 (One Health Pact). The funders had no role in study design, data collection and analysis, decision to publish or preparation of the manuscript.

## Acknowledgment

The authors would like to thank Thomas Langerak, Lisa Bauer, Feline Benavides, Lonneke Leijten, Daryl Geers Laurine Rijsbergen and Rik de Swart for technical assistance, Adam Meijer for providing the virus isolates, and Rory de Vries for helpful discussion.

## Supplementary Information

**S1 Fig.**
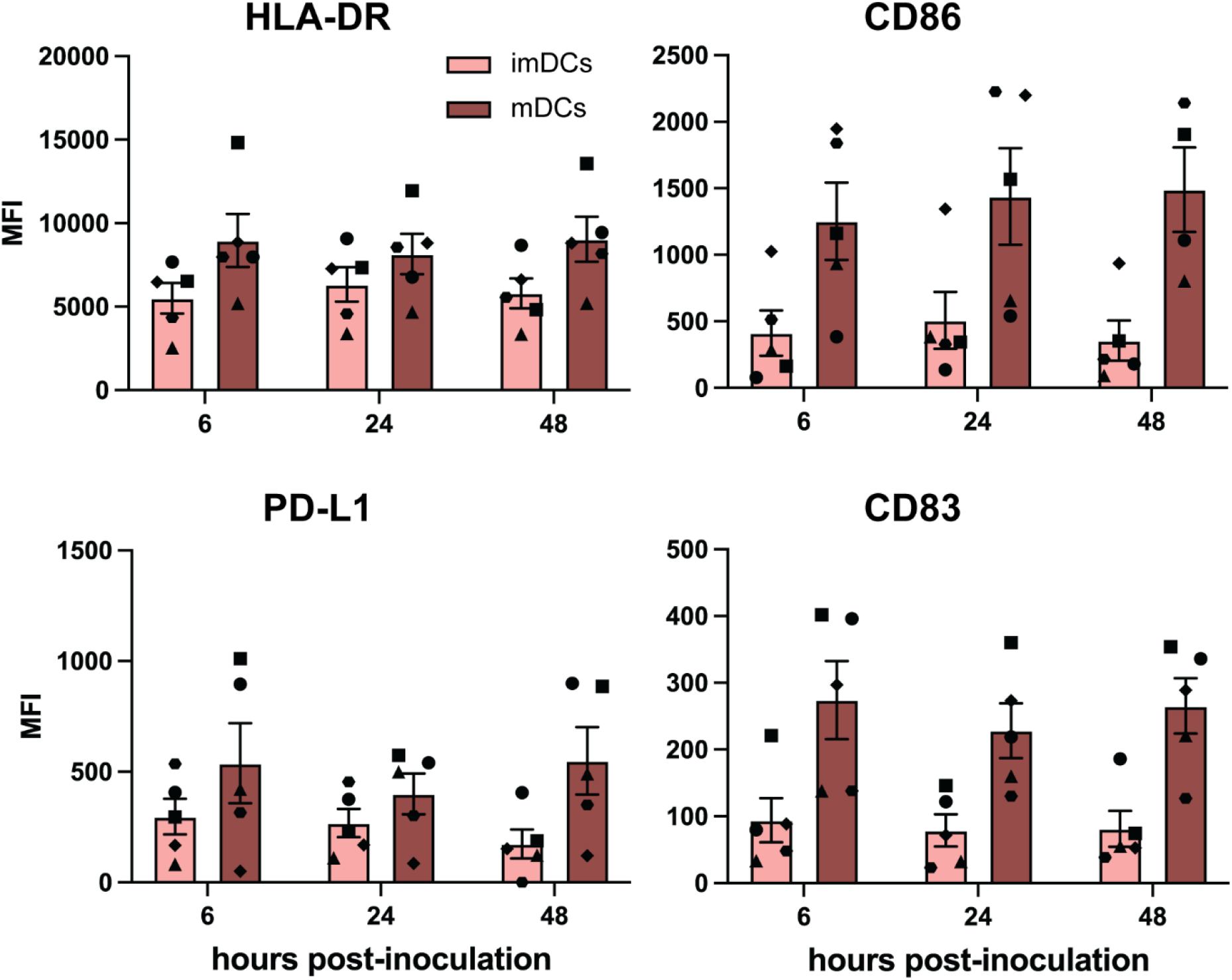
The expression of dendritic cell maturation markers in immature and mature dendritic cells (imDCs and mDCs). mDCs were defined by the upregulation of surface HLA-DR, CD86, PD-L1 and CD83 after treatment of monocyte-derived imDCs with lipopolysacharride. Each symbol represents one donor. Error bars denote standard error of mean. MFI: median fluorescence intensity.

**S2 Fig.**
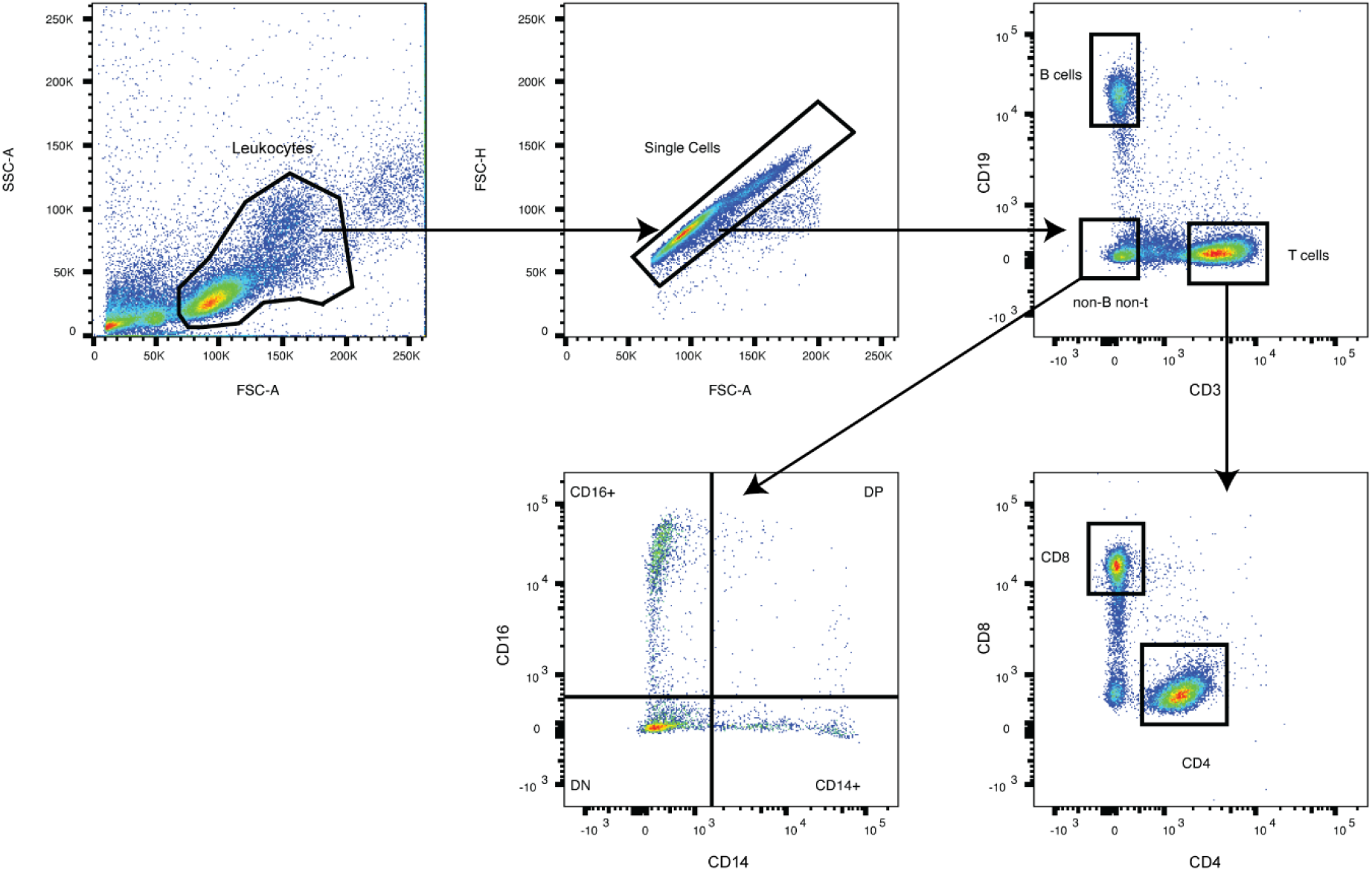
Representative gating strategy for peripheral blood mononuclear cell subpopulations. B cells were defined as CD3^-^CD19^+^ cells; CD4^+^T cells as CD3^+^CD19^-^CD4^+^CD8^-^ cells; CD8^+^T cells as CD3^+^CD19^-^CD4^-^CD8^+^ cells; and monocytes as CD3^-^CD19^-^CD14^-/+^CD16^-/+^ cells.

**S3 Fig.**
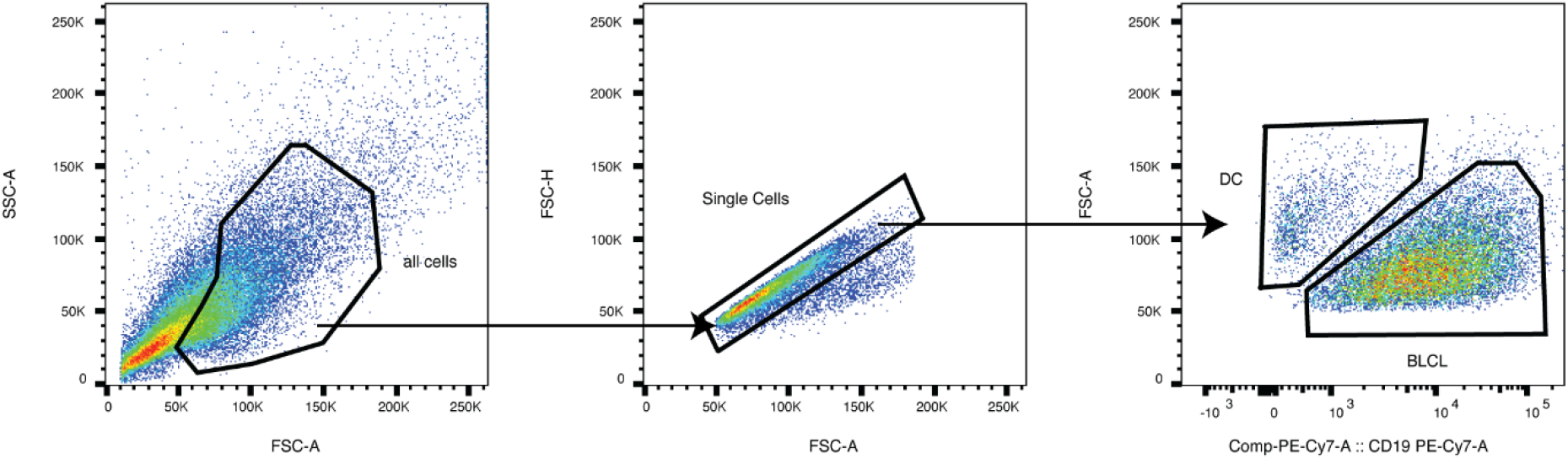
Representative gating strategy for EV-D68-inoculated immature dendritic cells (imDCs) co-cultured with autologous B-lymphoblastoid cell line (BLCL). imDCs were defined as CD19^-^ cells; BLCL were defined as CD19^+^ cells.

**S4 Fig.**
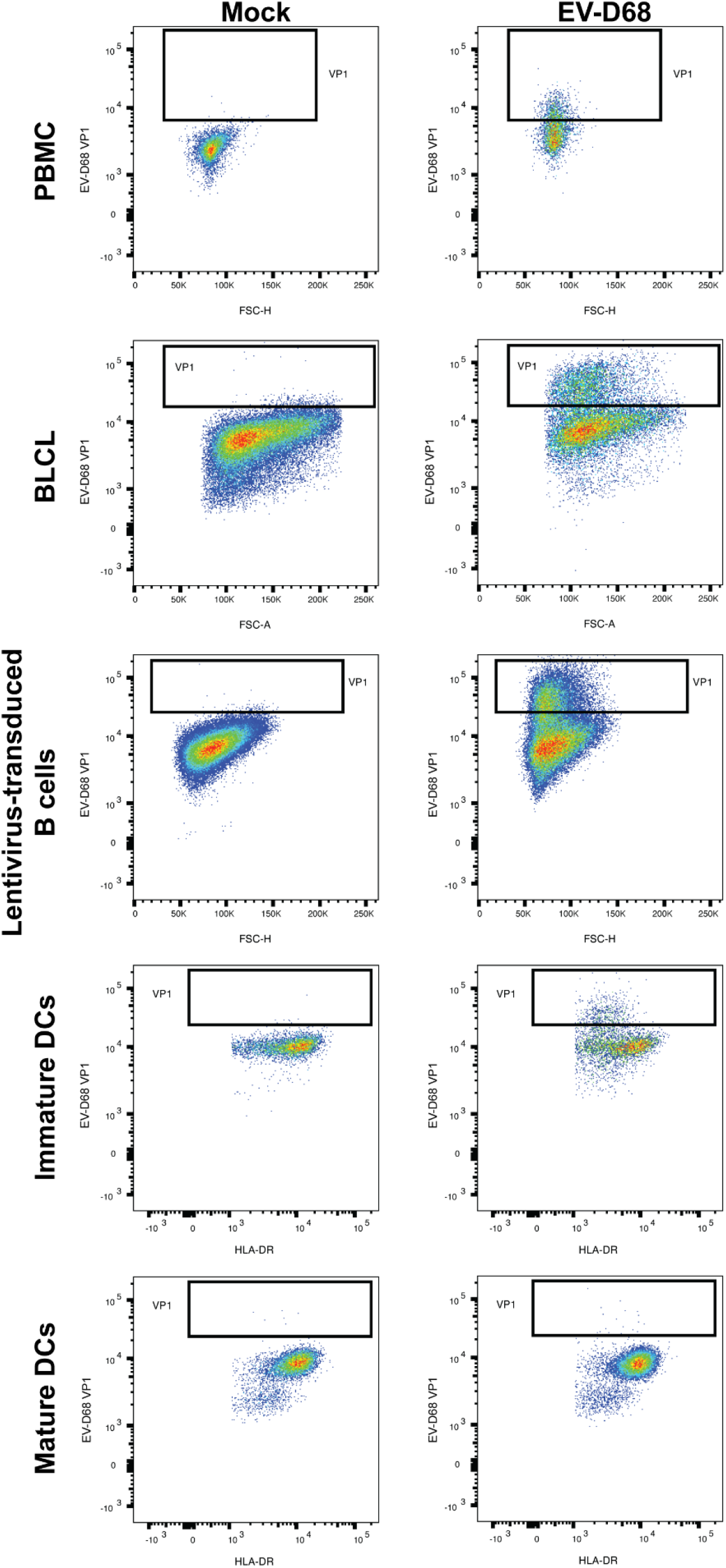
Representative gating strategies for EV-D68 VP1^+^ cells. PBMC: peripheral blood mononuclear cells; BLCL: B-lymphoblastoid cell line; DCs: dendritic cells.

**S5 Fig.**
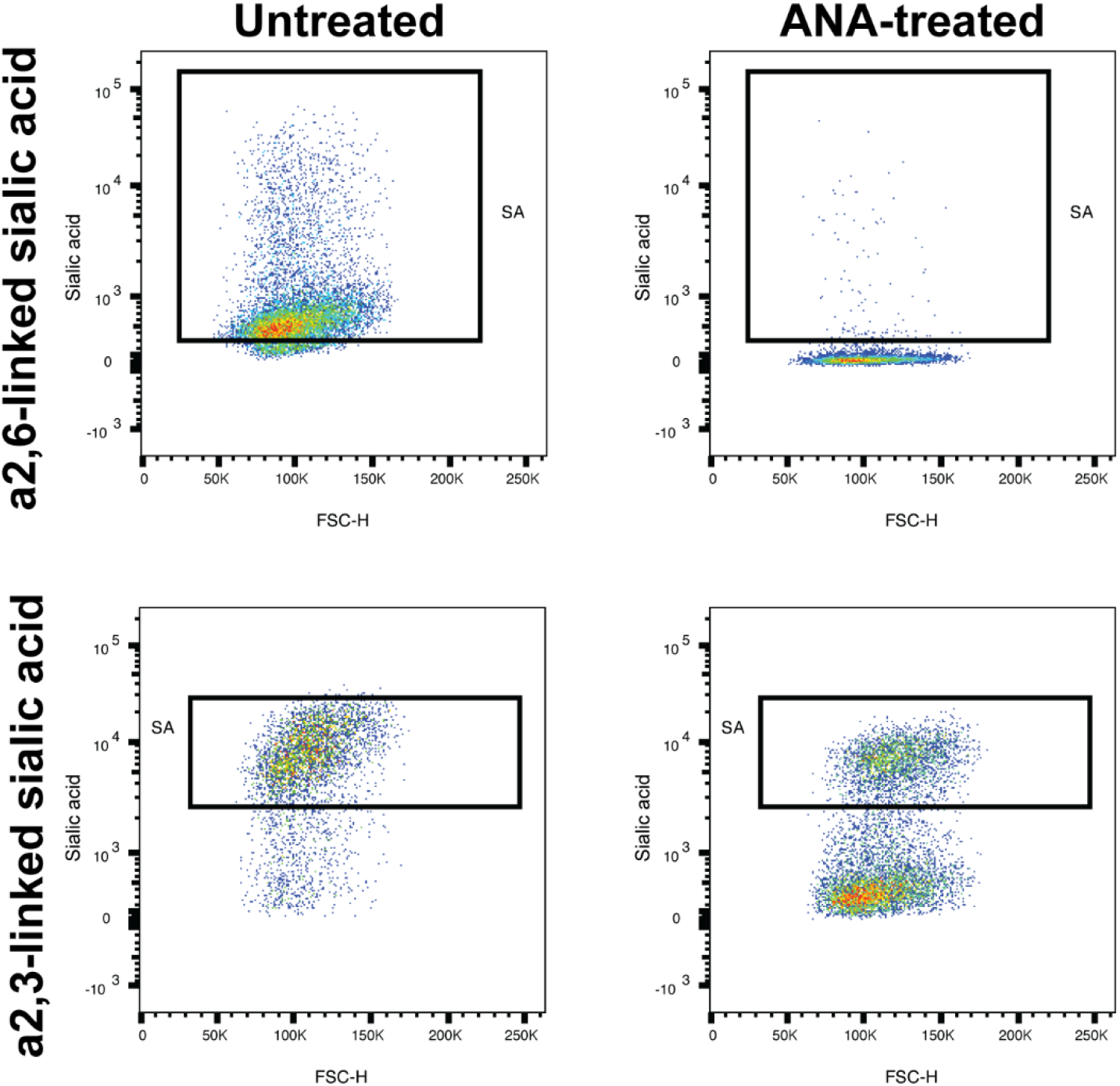
Representative gating strategy to determine percentages of α2,3- and α2,6-linked sialic acid^+^ (SAs^+^) BLCL. Percentages of SA^+^ cells were determined at 0 h post-inoculation. ANA: *Arthrobacter ureafaciens* neuraminidase; BLCL: B-lymphoblastoid cell line.

## Notes

### Competing Interest Statement

The authors have declared no competing interest.

